# Impact of a national tsetse control programme to eliminate Gambian sleeping sickness in Uganda: a spatio-temporal modelling study

**DOI:** 10.1101/2024.02.16.580671

**Authors:** Joshua Longbottom, Johan Esterhuizen, Andrew Hope, Mike J. Lehane, TN Clement Mangwiro, Albert Mugenyi, Sophie Dunkley, Richard Selby, Inaki Tirados, Steve J. Torr, Michelle C. Stanton

**Affiliations:** Department of Vector Biology, Liverpool School of Tropical Medicine, Liverpool, L3 5QA; Bindura University of Science Education, Bindura, Zimbabwe; Coordinating Office for Control of Trypanosomiasis in Uganda, Kampala, Uganda

**Keywords:** Tsetse, Spatio-temporal model, sleeping sickness, INLA, Uganda

## Abstract

**Introduction:** Tsetse flies (*Glossina*) transmit *Trypanosoma brucei gambiense* which causes gambiense human African trypanosomiasis (gHAT). As part of national efforts to eliminate gHAT as a public health problem, Uganda implemented a large-scale programme of deploying Tiny Targets, which comprise panels of insecticide-treated material which attract and kill tsetse. At its peak, the programme was the largest tsetse control operation in Africa. Here, we quantify the impact of Tiny Targets and environmental changes on the spatial and temporal patterns of tsetse abundance across north-western Uganda.

**Methods:** We leverage a 100-month longitudinal dataset detailing *Glossina fuscipes fuscipes* catches from monitoring traps between October 2010 and December 2019 within seven districts in north-western Uganda. We fitted a boosted regression tree model assessing environmental suitability which was used alongside Tiny Target data to fit a spatio-temporal geostatistical model predicting tsetse abundance across our study area (∼16,000 km^2^). We used the spatio-temporal model to quantify the impact of Tiny Targets and environmental changes on the distribution of tsetse, alongside metrics of uncertainty.

**Results:** Environmental suitability across the study area remained relatively constant over time, with suitability being driven largely by elevation and distance to rivers. By performing a counterfactual analysis using the fitted spatio-temporal geostatistical model we show that deployment of Tiny Targets across an area of 4000 km^2^ reduced the overall abundance of tsetse to low levels (median daily catch = 1.1 tsetse/trap, IQR = 0.85-1.28) with no spatial-temporal locations having high (>10 tsetse/trap/day) numbers of tsetse compared to 18% of locations for the counterfactual.

**Conclusions:** In Uganda, Tiny Targets reduced the abundance of *G. f. fuscipes* and maintained tsetse populations at low levels. Our model represents the first spatio-temporal model investigating the effects of a national tsetse control programme. The outputs provide important data for informing next steps for vector-control and surveillance.

**Key questions:** *What is already known on this topic?:* Small panels of insecticide-treated fabric, called Tiny Targets, are used to attract, and kill riverine tsetse, the vectors of *T. b. gambiense* which causes gambiense human African trypanosomiasis (gHAT). In large-scale (250-2000 km^2^) trials conducted in five countries, deployment of Tiny Targets reduced the densities of tsetse by between 60 and >90%.

*What this study adds:* We report an analysis of, and data from, a large-scale (∼4,000km^2^) national tsetse control programme, implemented in Uganda to eliminate gHAT as a public health problem. We found that Tiny Targets reduced tsetse abundance across the study period (2011-2019) and maintained densities at low (<1 tsetse/trap/day) levels. We produce maps which detail spatial variances in tsetse abundance in response to vector control.

*How this study might affect research, practice, or policy:* In 2022, Uganda received validation from the World Health Organisation (WHO) that it had eliminated gHAT as a public health problem. The large-scale deployment of Tiny Targets contributed to this achievement. Our findings provide evidence that Tiny Targets are an important intervention for other countries aiming to eliminate gHAT.

## Background

Human African trypanosomiasis (HAT), commonly called sleeping sickness, is caused by subspecies of *Trypanosoma brucei* transmitted by tsetse flies (*Glossina*). In West and Central Africa, gambiense HAT (gHAT) is caused by *T. b. gambiense* transmitted by riverine species of tsetse (e.g., *G. palpalis palpalis, G. fuscipes*). In East and Southern Africa, *T. b. rhodesiense* transmitted by savanna species of tsetse (e.g., *G. morsitans morsitans, G. pallidipes*) causes rhodesiense HAT (rHAT). Both diseases are fatal without medical intervention. Uganda is the only country where both forms of HAT occur [1].

The last major epidemic of HAT in Uganda occurred in the last ∼20 years of the twentieth century when political and economic upheavals disrupted national control programmes. Between 1990 and 1999, Uganda reported an average of 1,384 cases/year (range: 971-2,066) of gHAT and 516 cases/year (178-1,417) of rHAT [2]. Since then, numbers of both forms of HAT have declined. In the last five years for which data are available (2018-2022), there have been a total of four cases of gHAT and 13 cases of rHAT reported in Uganda. The dramatic decline in gHAT has been achieved through mass screening and treatment of human cases supported by the deployment of Tiny Targets to control tsetse [3]. For rHAT, the decline has been achieved through mass treatment of cattle with trypanocides and insecticides because in Uganda cattle are important reservoir hosts for *T. b. rhodesiense* [4] and cattle form the main source of a tsetse’s diet.

The achievements of Uganda over the past 20 years are part of a larger continental effort, led by the WHO, to eliminate gHAT as a public health problem by 2020 and eliminate transmission by 2030. Uganda’s achievement of the first goal was ratified by the WHO in May 2022 [5]. The second goal is defined as the “*reduction to zero of the incidence of infection in a defined geographical area, with minimal risk of reintroduction, as a result of deliberate efforts*”; this target involves 15 endemic countries by 2030 [6].

For the last decade, deployment of Tiny Targets has formed an important part of Uganda’s strategy to control gHAT. Tiny Targets are small panels composed of blue cloth (25 × 25 cm) flanked by a panel (25 × 25 cm) of black netting. The cloth and netting are impregnated with insecticide; tsetse are attracted visually to the target contact it and die [7]. In Uganda, Tiny Targets are deployed at a density of 20 targets per linear kilometre along the rivers and streams where riverine tsetse concentrate. This intervention reduces the density of tsetse populations by 60-99% [7-11]. Epidemiological models [12, 13] and empirical evidence [10, 11, 14] suggests that this reduction is sufficient to interrupt transmission. The very first trials of Tiny Targets were carried out in Uganda in 2011 [7] and from an initial trial covering ∼250 km^2^ the intervention grew to an operation of ∼4000 km^2^ across seven districts. At its peak, Uganda was implementing the largest national tsetse control operation in Africa. Tiny Targets are also making important contributions to the elimination of gHAT in Côte d’Ivoire [9], Chad [10], DRC [15] and Guinea [11].

The large-scale deployment of Tiny Targets in Uganda has been accompanied by an extensive monitoring programme comprising a network of entomological sentinel sites, where pyramidal tsetse traps are used to quantify the abundance of tsetse before and after targets were deployed [7, 8]. This monitoring programme has produced a decade of data on the distribution and abundance of tsetse in and near the places where Tiny Targets have been deployed in north-west Uganda.

Prior analyses of the impact of targets in Uganda [7, 8] and elsewhere [9-11, 15] have shown that the reductions in density varied between 55 and >99%. The causes of this variation are unknown but we hypothesize that the differences are due, in part, to underlying environmental factors. Previous estimates compared catches from individual sites before and after an intervention and could not consider what happened in places where we did not sample. The spatial and temporal scale of the monitoring data accompanying control operations in Uganda provides a unique opportunity to quantify the impact of a large-scale tsetse control programme and assess the relative contributions of Tiny Targets and environmental factors to the reduction in tsetse abundance throughout Uganda, in areas which were not measured empirically. To do this, we first developed temporally varying estimates of tsetse habitat suitability within north-western Uganda, using pre-intervention entomological survey data, remotely sensed environmental data and a species distribution model. Suitability outputs were then combined with data from subsequent post-intervention surveys to quantify the impact of both environmental change and vector control on the abundance of *Glossina fuscipes fuscipes*, the main g-HAT vector in Uganda, through a spatio-temporal geostatistical modelling approach.

## Methods

### Study area

Trapping was performed to quantify the impact of Tiny Targets on the abundance of tsetse [7, 16]. Between October 2010 and December 2019, pyramidal traps [17] were deployed within seven districts in north-western Uganda to monitor the abundance of *G. f. fuscipes* [7, 18]. Traps were deployed for 1-4 consecutive days (median 2 days), with tsetse collected and counted at 24-hour intervals [7, 19]. Monitoring and control activities were scaled-up in phases according to need and available funding, therefore initial deployment of traps and targets varied between and within districts. The year in which intervention was initiated in each district is displayed in Figure 1, utilising watersheds as a nominal metric of coverage [14] – further detail regarding survey dates and distribution are provided within Appendix: Supplementary Methods.

**Figure 1:**
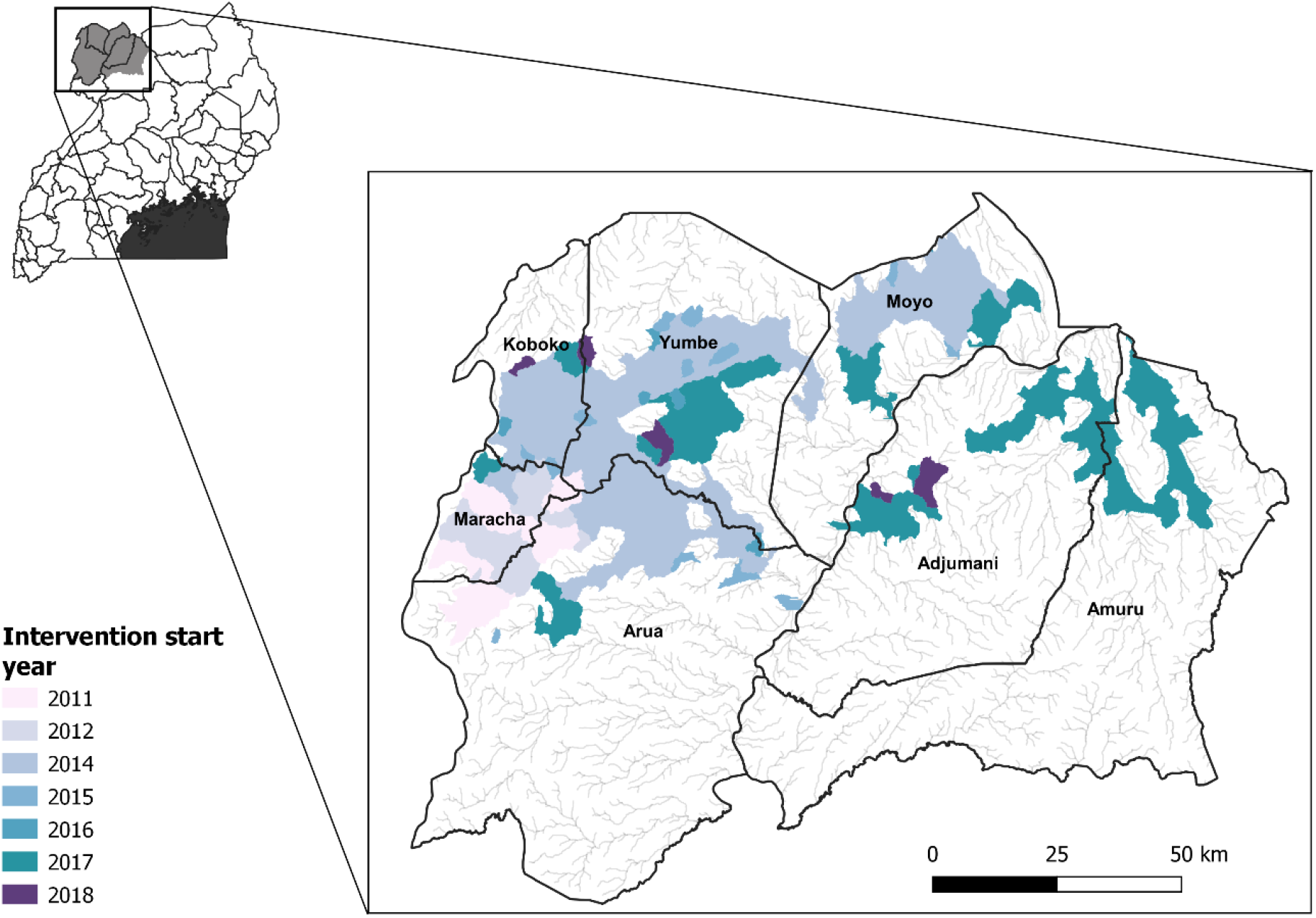
Districts within north-western Uganda in which tsetse monitoring was performed, alongside watersheds controlled by Tiny Targets. The seven districts which form the basis of this analysis cover a total of ∼16,419km^2^, combined coverage of intervention areas is ∼4,000 km^2^. Colours represent the years in which the Tiny Target intervention was first rolled out within each district, constructed using data from [14]. Targets were replaced every six months. Map produced using QGIS 3.16.5 [20].

### Identification of intervention areas

Tiny Targets are along riverbanks within the intervention area twice a year. They are deployed at 100m intervals along each bank, i.e., 20 targets per kilometre of river, with their location recorded using global positioning systems (GPSs). We generated a 30m × 30m resolution grid for the entire study area and the distance from the centre of each grid cell to the nearest Tiny Target was calculated per deployment period. We assumed that targets were effective for six months following their deployment (Appendix: Table 1) based on other work [7]. These distance surfaces were used to produce a categorical variable classifying each gridded pixel as (1) within 500m of a target, (2) >500m but <5000m or (3) >5000m from a target [15]. Henceforth, the three categories are termed Inside, Edge and Outside respectively.

**Table 1:** Description and source of covariates used within the presence-absence modelling framework.

| *Temporal resolution* | *Covariate (unit)* | *Rationale (reference)* | *Source* |
| --- | --- | --- | --- |
| *Synoptic* | Elevation (meters) | [23, 24] | Shuttle Radar Topography Mission (SRTM) [27] |
|  | Distance to rivers (meters) | [26] | Derived from Shuttle Radar Topography Mission (SRTM) elevation data [27] |
|  | Slope (percentage gain) |  | Derived from Shuttle Radar Topography Mission (SRTM) elevation data [27] |
| *Annual (2011-2019)*<br>*[Dry season]* | Land surface temperature (LST) day (mean) (°Celsius) | [21, 22] | Derived from Landsat 5 [28]<br>Derived from Landsat 8 [29] |
|  | Normalised difference vegetation index (NDVI) (-1 – 1) | [25] | Derived from Landsat 5 [28]<br>Derived from Landsat 8 [29] |
|  | Proportion of vegetation (PVI) (0 – 1) | [25] | Derived from Landsat 5 [28]<br>Derived from Landsat 8 [29] |

### Tsetse data for model

Geographic locations of monitoring traps were recorded using GPSs alongside additional variables outlined in Appendix: Table 2. From collected records, we produced two separate datasets: one for use in a species distribution model (SDM) predicting habitat suitability, and another for use in a geostatistical spatio-temporal modelling framework predicting tsetse abundance over time. The main differences between the data requirements for the two models were that the spatio-temporal geostatistical model was fitted to count data of tsetse from all traps, and the SDM used presence-absence data with observations being limited to traps considered to be unaffected by the intervention, i.e., operated before any intervention or ≥5km from a Tiny Target. As the geographical extent of the intervention increased, some traps classed initially as being ‘non-intervention’ transitioned to an ‘intervention’ status.

**Table 2.** Posterior mean estimates and credible intervals (CrI), alongside rate ratio estimates for the best fitting model (Model 10, Supplementary Table 6). Blank (dashed) rows represent the reference category used with each interaction term. LTT = Linear temporal trend, RF = Gaussian random field, *CrI* = Credible interval.

| Variable | Mean | 2.5% CrI | 50% CrI | 97.5% CrI | Rate ratio (95% CrI) |
| --- | --- | --- | --- | --- | --- |
| Suitability | 0.089 | -0.079 | 0.089 | 0.257 | 1.09 (0.92, 1.29) |
| Inside intervention area (<500m) | 0.420 | -30.647 | 0.420 | 31.461 | 1.52 (4.90e<sup>-14</sup>, 4.61e<sup>13</sup>) |
| Edge of intervention area (>500m, ≤5000m) | 0.769 | -30.299 | 0.768 | 31.810 | 2.16 (6.94e<sup>-14</sup>, 6.53e<sup>13</sup>) |
| Outside of intervention area (>5000m) | -0.036 | -31.104 | -0.036 | 31.006 | 0.96 (3.30e<sup>-14</sup>, 2.85e<sup>13</sup>) |
| Season: Dry | - | - | - | - | - |
| Season: Wet | 0.011 | -0.070 | 0.011 | 0.092 | 1.01 (0.93, 1.10) |
| Linear temporal trend (LTT) | -0.025 | -0.027 | -0.025 | -0.023 | 0.98 (0.97, 0.98) |
| Suitability*Inside of intervention area | - | - | - | - | - |
| Suitability*Edge of intervention area | 0.387 | 0.035 | 0.388 | 0.734 | 1.61 (0.96, 2.69) |
| Suitability*Outside of intervention area | 1.463 | 1.056 | 1.464 | 1.867 | 4.72 (2.66, 8.36) |
| Wet season*Inside of intervention area | - | - | - | - | - |
| Wet season*Edge of intervention area | 0.044 | -0.135 | 0.044 | 0.223 | 1.06 (0.81, 1.37) |
| Wet season*Outside of intervention area | 0.269 | 0.098 | 0.269 | 0.439 | 1.32 (1.03, 1.70) |
| LTT*Inside of intervention area | - | - | - | - | - |
| LTT*Edge of intervention area | -0.009 | -0.012 | -0.009 | -0.005 | 0.97 (0.96, 0.97) |
| LTT*Outside of intervention area | -0.009 | -0.013 | -0.009 | -0.005 | 0.97 (0.96, 0.97) |
| Spatial range of the RF: $\rho$ | 0.097 | 0.095 | 0.097 | 0.715 | - |
| Marginal standard deviation of the RF: $\sigma^2$ | 1.597 | 1.578 | 1.597 | 1.616 | - |
| Size for nbinoimial zero-inflated observations: $\alpha$ | 0.704 | 0.699 | 0.704 | 0.710 | - |
| Zero-probability parameter for zero-inflated nbinoimial: $\pi$ | 0.039 | 0.039 | 0.039 | 0.040 | - |

### Assembling explanatory variables

Gridded surfaces for temperature, elevation and vegetation were assembled for the seven districts (extent shown in Figure 1, covariates summarised in Table 1). These variables have been shown previously to influence the distribution of tsetse [21-25]. Covariates were generated from remotely sensed satellite imagery collected at a spatial resolution of 30 × 30m during the dry season – December-February. Cloud coverage and clear scene availability affected our capability to collate imagery for other times of year. Where available, temporally varying covariates were collated annually. Non-temporally varying covariates (elevation, distance to rivers, slope) were included as synoptic surfaces (Table 1). To account for tsetse dispersal [26], a buffer with a radius of 150m was used to produce a smoothed mean covariate derived from averaging all 30m cells across 300m of the true sample location. An overview of the covariate production process is provided in the Appendix, Supplementary Methods.

### Species distribution model

To estimate tsetse densities in locations where no sampling was performed, we produced annual estimates of habitat suitability using a presence-absence species distribution model (SDM). SDMs predict the distribution of a species across a landscape [30] by combining information on species occurrence with environmental variables (covariates) at the same location [31]. Using the species occurrence dataset and covariates detailed above, we constructed a presence-absence boosted regression tree (BRT) model using the ‘*caret*’ package within R (version 3.5.1) [32, 33].

BRTs are a machine learning algorithm which combine both regression trees and boosting (iteratively combining a group of simple models) to build a linear combination of many trees [34], and have been used to predict the distributions of a number of diseases and disease vectors [15, 35-37]. The BRT method models a suitability index from 0 to 1 for tsetse based on the values of environmental covariates at the locations corresponding to presence-absence inputs [30]. In this instance, ‘*presence*’ and ‘absence’ records refer to sampling locations where tsetse were caught or not. The absence of tsetse from a monitoring trap may reflect the true absence of tsetse or that tsetse were present but the trap failed to catch any. Accordingly, ‘*absence*’ records are commonly referred to as ‘*background points*’ and serve the purpose of exposing the model to locations where the species is presumed to be absent [38]. Further information regarding the covariates used within the model, model fitting, and methods of model evaluation can be found in the Appendix: Supplementary Methods.

### Spatio-temporal model

To evaluate the impact of Tiny Targets and environmental variables on tsetse, a geostatistical spatio-temporal model was constructed using catch data. Data for this model included repeat catches from sites within the same year. Prior to constructing the spatio-temporal model, a series of exploratory plots and analyses were performed. An empirical variogram was constructed to test for spatial autocorrelation and to obtain starting parameter values for use within the model; this variogram was fit using the ‘*PrevMap*’ R package [39]. A Pearson’s Chi^2^ test was performed to determine the level of dispersion within the data using the ‘*msme*’ R package [40]. Given the high number of zero catches, our *a priori* assumption was a high level of overdispersion. This was confirmed (dispersion = 4.79), and therefore a negative binomial distribution was most appropriate for modelling [41]. As trapping success is highly variable, and zeros may arise due to either the true absence of the species, or due to trapping failure, we opted to model excess zeros independently through use a zero-inflated negative binomial (ZINB) model.

The underlying statistical model was a spatially and temporally explicit hierarchical generalised linear regression model for ZINB data, using the log link function and a type 1 likelihood. A type 1 likelihood accounts for two different types of zeros within the dataset: *structural* or true zeros which represent the true absence of tsetse in a location, and *sampling* zeros, where a zero is recorded as a reflection of chance [42]. Further information regarding model construction, including a full model description, can be found in the Appendix: Supplementary Methods.

### Model fitting and validation

Models were fit through integrated nested Laplace approximations (INLA) and a stochastic partial differential equation (SPDE) representation of the Gaussian-Markov random field (GMRF) approximation to the Gaussian process model, based on a Matérn covariance function, using the R INLA package [43]. To evaluate the significance of fixed and random effects on tsetse abundance, an iterative process was performed where varying combinations of fixed and random effects were used within models to identify the optimal model construction.

*A priori*, we hypothesised that the effect *‘suitability’*, the output of the BRT environmental suitability model, would be positively associated with the catch of tsetse. We hypothesised further that the ‘*intervention*’ effect, i.e., proximity to Tiny Targets, would have a negative association on tsetse abundance. An interaction term between ‘suitability’ and ‘intervention’ was also included in the fitting process. Temporal effects were included in the form of *‘season’*, a categorical ‘wet’ and ‘dry’ variable, to investigate temporal changes in abundance and through the inclusion of an autoregressive process of order 1, i.e., AR(1). It was hypothesised that there will be seasonal changes in abundance, with a higher abundance of tsetse being observed in the wet season [7, 44].

Measures of the goodness of fit for the spatio-temporal model were obtained using the deviance information criterion (DIC). The DIC is a Bayesian generalization of the Akaike information criterion (AIC), where models are penalised by their deviance and the number of parameters included [45]. Using the variables identified from the model with the lowest DIC, we fitted a separable geostatistical spatio-temporal model described fully in Appendix: Supplementary Methods. The INLA approach does not allow for the combined fitting of a regression model for the zero-inflation probability of the zero-inflated model, therefore an additional function (‘pred.zinb’) was defined to apply the zero-inflation probability to a posterior sample (1000 draws) of non-zero inflated data derived from the negative binomial model [46]. Model validation was performed using a spatial leave-one-out cross-validation (SLOO-CV) approach, based on an adaptation of methods described in [47, 48], and described further within the Appendix: Supplementary Methods. Validation statistics included assessing the correlation between the predicted and observed tsetse densities through summaries of the root-mean-square error (RMSE) and the mean absolute error (MAE) [49].

Posterior predictive distributions (1000 draws) were simulated for each 30 × 30m cell to determine the probability of tsetse catches exceeding predefined abundance categories defined as *low*, 0-1 flies, *medium*, >1 - 10 flies, and *high*, >10 flies. These categories were determined by discussion with field entomologists regarding policy-relevant values. The number of draws within each category was used to produce probabilities for each cell, low (*p*_*L*_), medium (*p*_*M*_), and high (*p*_*H*_) respectively, and each category was assigned a predictive score using the log-odds. Following Lowe *et al*. [50], we used the receiver operating characteristic (ROC) to define optimal probability thresholds for assigning a final category to each cell by comparing the predictive score with the observed class, using the ‘ROCit’ R package [51]. The probabilistic results were mapped using a ternary plotting technique [52], and the ‘*tricolore*’ R package [53] to visualise category certainty. Within the maps, the predicted category for each cell was expressed as a colour determined by a combination of the three probabilities assigned, with colour saturation used to indicate the associated certainty. Maps of the final category per cell were produced using QGIS and threshold values obtained from the ROC curves.

### Counterfactual analysis

To estimate the relative contribution of Tiny Targets to changes in tsetse abundance, a counterfactual analysis was performed using a 50% random sample of the longitudinal trapping data, i.e., 4180 trap-month records. Using the optimal spatio-temporal model, all covariate values were held fixed except for the categorical intervention variable. The predicted mean flies per trap day was then compared for two predictions, where the value assigned to the intervention variable was changed:

1. 50% of records were assigned the intervention category ‘inside’ for the purpose of prediction and estimates of the mean daily catch of tsetse were generated for these locations.
2. In a different model run, the same 50% of records were assigned the intervention category ‘outside’ and estimates of the mean daily catch were generated for these locations.

To compare the abundances predicted by the original and counterfactual models, we quantified changes in the frequency of catches in the low, medium and high catch categories.

## Results

### Tsetse occurrence data

The dataset comprised 31,426 records from 569 locations sampled between October 2010 and December 2019 (Figure 2). A total of 52,544 tsetse were captured over 31,553 trapping days (mean 1.67 flies/trap/day across all locations) (Appendix: Table 3).

**Figure 2.**
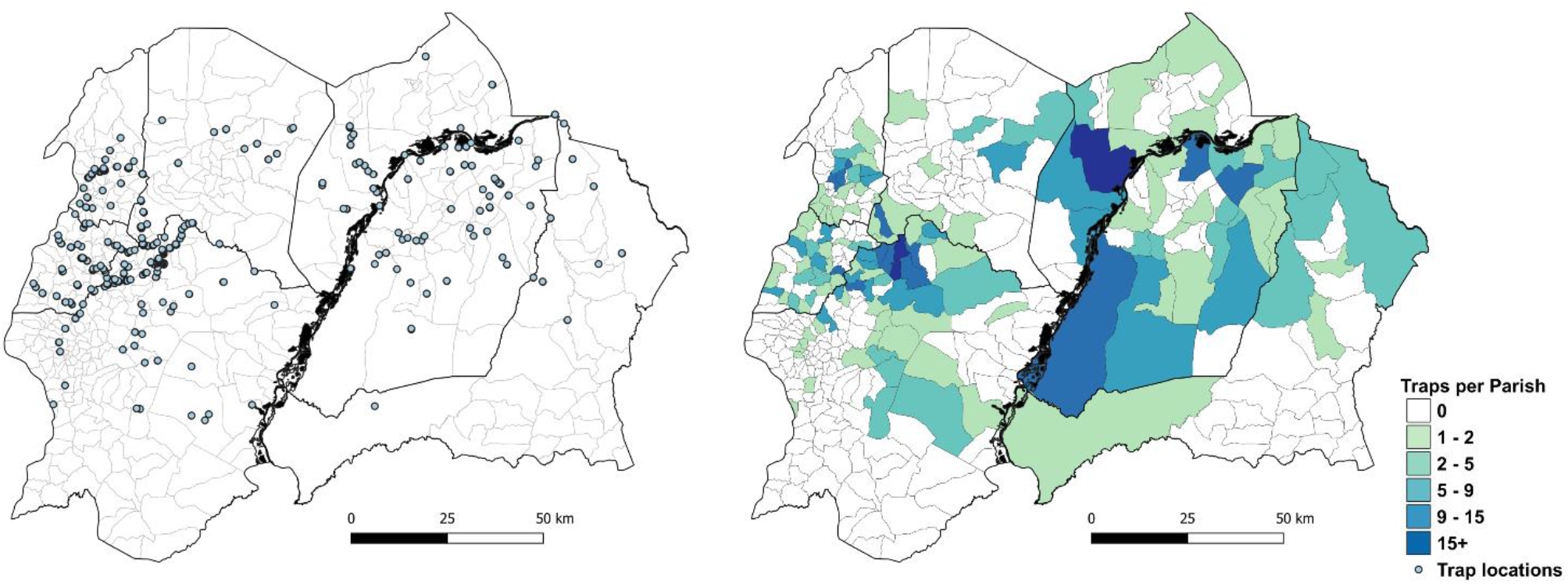
Location (left) and number of tsetse traps per parish (right), north-western Uganda. Parish administrative boundaries obtained from www.GADM.org. Map produced using QGIS 3.16.5 [20].

After spatial and temporal aggregation to retain one record per 30m × 30m cell per year (presence at one time point replaces absence at another time within the same year), 538 unique location-year records situated outside the intervention area remained: 376 presence and 162 absence. The number of records per year is provided as Appendix: Table 4. We sampled predominantly where we presumed tsetse to be present and so locations outside the intervention area reporting absences are relatively few.

### Habitat suitability maps

A BRT model was fitted to presence-absence data from 2010-2019 obtained outside the intervention area. Optimal values of the BRT model parameters, based on minimising the Brier score, were number of trees = 350, interaction depth = 29 and shrinkage = 0.1. The evaluation measures for the resulting optimal BRT are as follows: AUC = 0.81 and Brier score = 0.21, representing a moderate model fit. The specificity and sensitivity of the model were 0.86 and 0.59 respectfully, indicating a greater ability to correctly identify absence records (high specificity) and a greater error when predicting presence locations (low sensitivity). The BRT model may be calibrated to favour optimising either sensitivity or specificity or calibrated to equally prioritise both. For this study, we used default settings for balancing sensitivity and specificity.

The relative importance of each of the environmental variables included, with respect to their contribution to the final BRT, is presented in Appendix: Table 5. Elevation (m), NDVI and distance to rivers (m) were equally important contributors (19.14%, 18.57%, 18.47% respectively), to variation in suitability. Using the fitted BRT model, predictions of habitat suitability for tsetse were made for each 30m x 30m cell for eight years: 2010, 2013-2019. Maps showing suitability for the years 2010 and 2019 are presented in Figure 3; maps for other years are provided as Appendix: Figure 1. Areas of high suitability follow rivers, vegetated and high-elevation areas neighbouring the Albert Nile and parts of Amuru and Adjumani districts. Spatial trends are visible across years, and for parts of Amuru and Adjumani districts predicted suitability was greater in 2019 than 2010 ignoring the impact of targets, however, this may be an artefact of the low sensitivity of the model (0.59).

**Figure 3.**
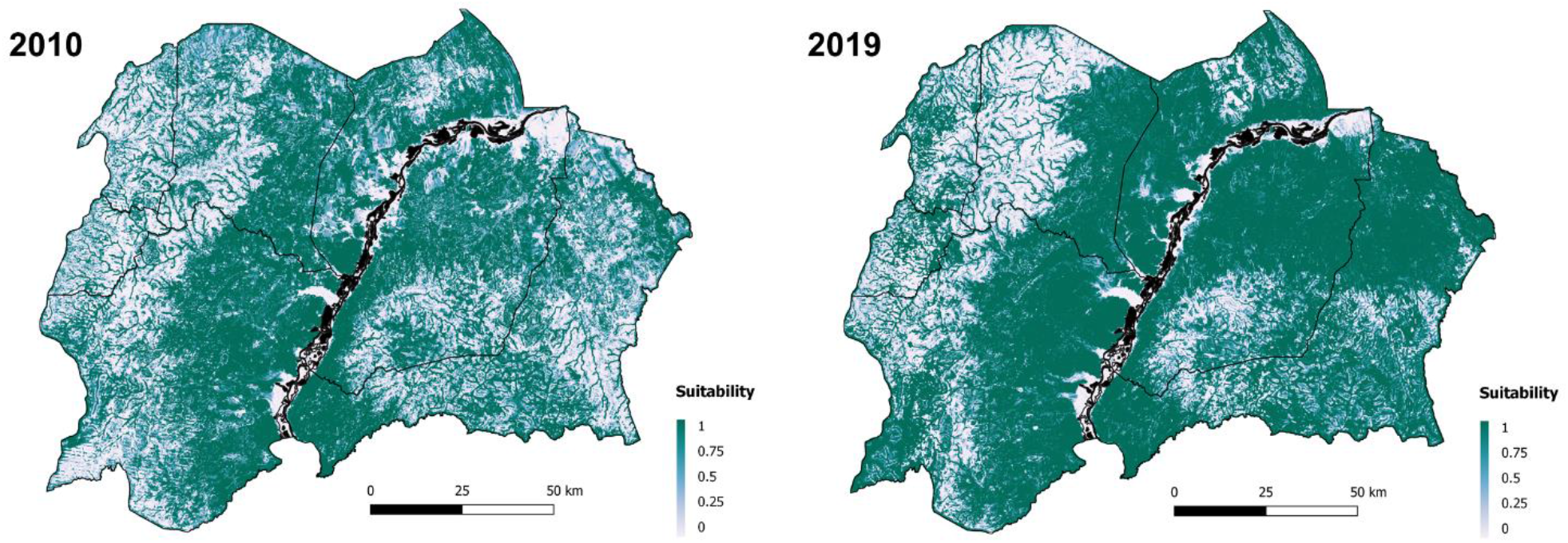
Predicted habitat suitability for *Glossina f. fuscipes* within north-western Uganda, for 2010 and 2019. Dark green locations indicate areas of higher environmental suitability; whiter areas indicate areas of lower environmental suitability. Map created using QGIS 3.16.5 [20].

### Spatio-temporal Model

After collating all sampling records, a 100-month continuous longitudinal dataset consisting of 416 sites was produced for north-western Uganda. The series consisted of records from September 2011 to December 2019 and included 8360 trap-month combinations (total flies [count] reported at a trap, for a specific month). These data formed the basis of the spatio-temporal model.

To identify which fixed and random effects optimise the performance of the spatio-temporal model, a range of ZINB generalised linear geostatistical models (GLGMs) were fitted to the series data, varying the fixed and/or random effects across models. A list of considered models, alongside their corresponding evaluation metrics (DIC, WAIC and CPO) is provided as Appendix: Table 6. The optimal ZINB had an DIC of 35538, compared with a median and maximum DIC of 36100 and 37064, respectively (Appendix: Table 6).

The equation for the final model is as follows:

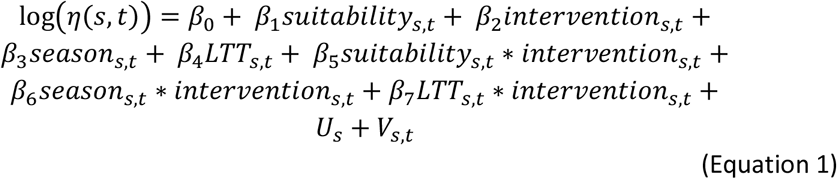

*β*_*n*_, *n = 1, …, 7* represents the coefficients for each covariate associated with observations at location *s* at time *t*. The abbreviation *LTT* refers to a linear temporal trend, representing the sequential month in the continuous time series. *U*_*s*_ represents the spatially uncorrelated random effect (*site*_*ID*_), and *V*_*s,t*_ represents the spatially and temporally structured random effects fully defined in Appendix: Supplementary Methods.

Table 2 displays the posterior mean estimates and 95% Bayesian credible intervals (*CrI*) for the effects included within the optimal spatio-temporal model, fit to all observed locations and time periods (100-month series). Posterior distributions and *CrI* are visualised as Appendix: Figure 2. Starting parameter and hyperparameter values for priors used within the model are provided within Appendix: Table 7.

We performed a SLOO-CV using the optimal spatio-temporal model configuration. In total, 415 separate sub-models were produced (*K − 1*). Each sub-model was fitted to data excluding one randomly selected trap, and all traps within a radius of 0.097 decimal degrees from that trap. This radius was equivalent to ∼12.8km, the value defined by the posterior range of spatial autocorrelation (*ρ*) identified within the optimal model. The zero-inflation probability (*π* = 0.039, 95% *CrI* = 0.039, 0.040) was applied to a posterior sample consisting of 1000 draws of non-zero inflated data within each model, and the mean predicted values across all 1000 draws for the excluded data were compared to the observed values. Resulting validation statistics include a RMSE of 8.021 and MAE of 6.048, lower than another published model from the region (RMSE = 15.2)[16].

Through the conversion of posterior mean values into rate ratios, we determined the mean effect of each variable on the predictions. Habitat suitability is significantly and positively associated with fly catches outside of the intervention area (Rate Ratio [*RR*] = 4.72, 95% *CrI* = 2.66, 8.36). This effect weakens at the edge of intervention areas (*RR* = 1.61, 95% *CrI* = 0.96, 2.69) and inside the intervention areas (*RR* = 1.09, 95% *CrI* = 0.92, 1.29) (Table 2). Other significant predictors include a linear temporal trend (significant negative effect, *RR* = 0.98, 95% *CrI* = 0.97, 0.98), and interactions between (1) season (wet) and intervention (significant positive effect, edge *RR* = 1.06, 95% *CrI* = 0.81, 1.37; outside *RR* = 1.32, 95% *CrI* = 1.03, 1.70), implying higher abundance of tsetse within the wet season and on the edge and outside of intervention areas compared to the dry season reference class, and (2) intervention and the linear temporal trend (significant negative effect, *RR* = 0.97, 95% *CrI* = 0.96, 0.97) (Table 2, Appendix: Figure 2).

For traps inside the intervention areas, the counterfactual increased the median daily catch from 1.1 (IQR = 0.93-1.28) to 2.3 (IQR = 1.79-7.14) tsetse/trap. Conversely, for traps outside the intervention area, the counterfactual reduced the median daily catch from 3.8 (IQR = 0.77-14.17) to 1.0 (IQR = 0.94-1.28) tsetse/trap. The counterfactuals had marked effects on the frequency distribution of catches in the high categories (Figure 4). For traps inside the intervention area, none were predicted to have mean daily catches of >10 tsetse/trap whereas for the counterfactual, 18.0% were in this category. Conversely, for traps outside the intervention area, 33.8% were predicted to have high catches compared to none for the counterfactual.

**Figure 4.**
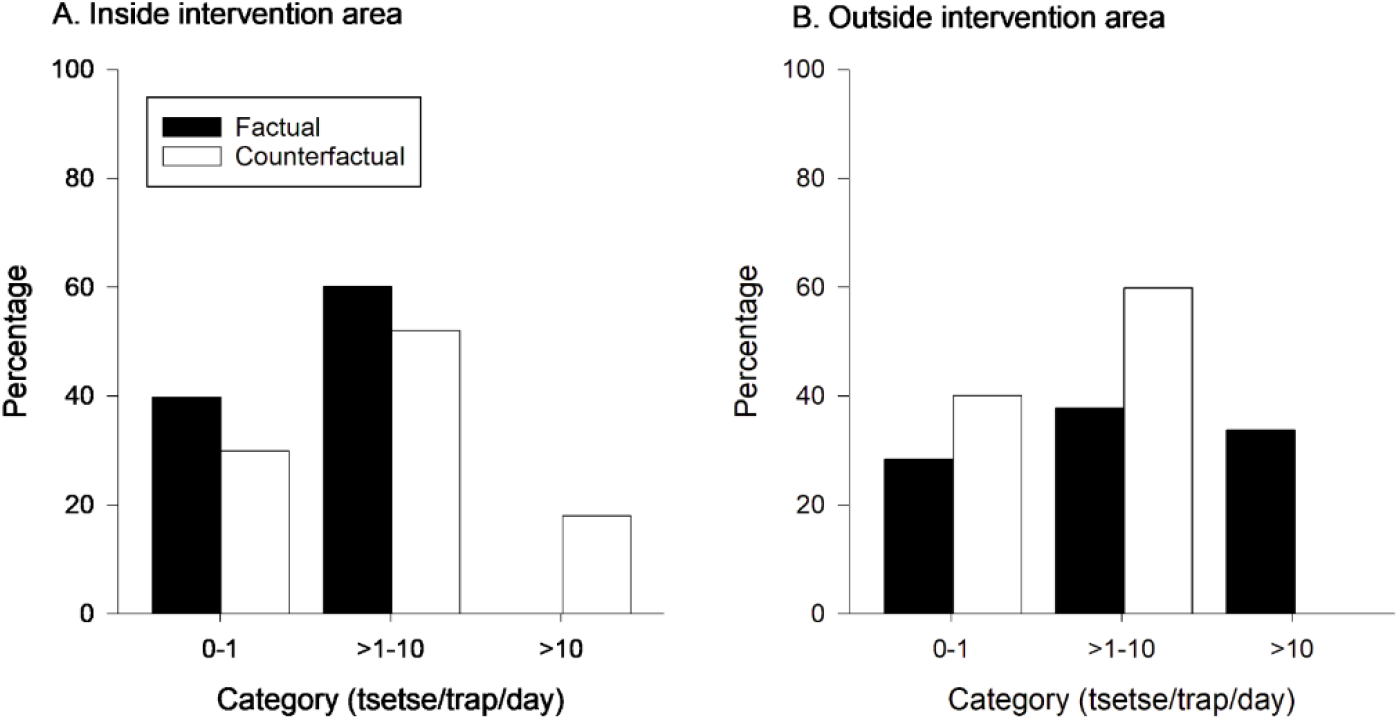
Frequency of catches (tsetse/trap/day) in low (0-1), medium (>1-10) and high (>10) categories for the factual and counterfactual models of tsetse abundance.

To determine threshold values for converting predictions into categorical estimates, a sample of 1000 draws were taken from the posterior distribution of the fitted model for each gridded cell. The probability of the prediction belonging to low, medium, and high fly categories was produced (*p*_*L*_, *p*_*M*_ and *p*_*H*_ respectively). These probabilities were compared with the true category from observed data, and ROC curves were generated (Appendix: Figure 3). The model was able to distinguish between ‘low’ and ‘high’ categories with high accuracy, with AUC values of 0.83 and 0.91. In the operational setting, identifying these extremes will help prioritise area for control and/or monitoring. The ability to correctly identify the ‘medium’ category (between 1-10 flies) was lower than the other two, with an AUC value of 0.7 (Appendix: Figure 3). Using the ROC curves, threshold values were obtained for assigning predictions to a specific category. If *p*_*L*_ ≥ 0.44, a cell was assigned the category ‘low’ abundance, if *p*_*L*_ < 0.44 and *p*_*M*_ ≥ 0.468, a cell was assigned the category ‘medium’ abundance, if *p*_*L*_ < 0.44 and *p*_*M*_ < 0.468, a cell was assigned the category ‘high’ abundance.

Predictions of tsetse abundance were produced for four periods: February 2012, 2015, 2017 and 2019, representing in turn (i) first trials of Tiny Targets [7], (ii) first expansion from two to five districts [8], (iii) second expansion to cover seven districts and (iv) time of maximum coverage (2019). The outputs are visualised as ternary maps, displaying the assigned abundance category (low, medium, and high) and associated certainty per gridded cell (Figure 5). Plots showing the relative abundance of tsetse for each period are provided as Appendix: Figure 4.

**Figure 5.**
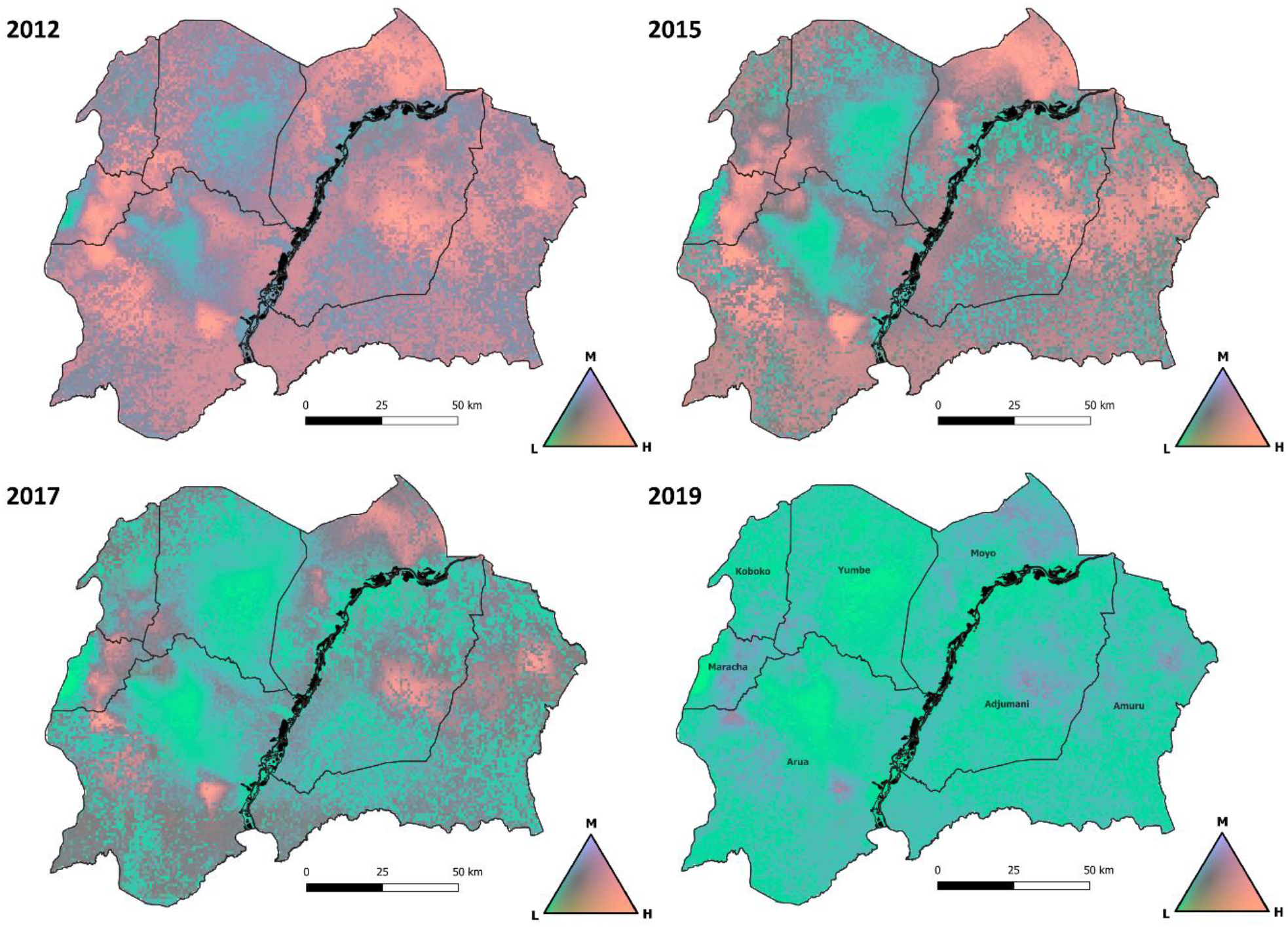
Comparison of categorised (low, medium, and high) *G. f. fuscipes* abundance during four time periods. The prediction period relates to February of each year (2012, 2015, 2017 and 2019). The continuous colour palette portrays the probabilities assigned to low, medium and high-abundance categories, with the low category representing 0 flies, medium 1-10 flies and high >10 flies. The greater the vibrancy, the more certain the prediction. Vibrant pink represents a high probability of a high-abundance of tsetse, vibrant green represents a high probability of low-abundance.

Comparing the categorical predictions from 2012 with those for 2019 (Figure 5) highlights a striking reduction in tsetse abundance over time. Generally, many ‘high’ abundance areas transition to areas of ‘low’ abundance between the two periods, starting with Yumbe district (2015) and expanding to areas of Adjumani, Arua and Amuru in 2017. The overall relative distribution and abundance of tsetse, however, does not appear to change across years (Appendix: Figure 4) despite the overall reductions in absolute abundance (Appendix: Table 8). Persistent tsetse populations, albeit with a lower abundance, can be seen in northwest and eastern Arua, Maracha, central Adjumani and north-eastern Amuru (Figure 5 & Appendix: Figure 4). Several of the relatively high abundance areas, such as central Adjumani and eastern Arua, are in places outside the 2019 intervention area (Appendix: Figure 4). Maps representing the categorical prediction after applying the threshold values determined by the ROC curves are given as Appendix: Figure 5; these maps can be used to inform additional monitoring and tsetse control operations.

## Discussion

We used a 100-month series of catches of tsetse from traps deployed at 416 sites across North-Western Uganda to produce species distribution and spatio-temporal models of the abundance of *G. f. fuscipes*, an important vector of sleeping sickness in Uganda and neighbouring countries (South Sudan, Democratic Republic of Congo). The species distribution model showed that the presence of tsetse was correlated negatively with elevation and positively with NDVI and proximity to rivers, in accordance with previous studies [15, 16]. Whilst temporal variation in habitat suitability occurs over the study period, few locations show trends of decreased suitability (areas in Maracha, Koboko, Arua and Yumbe when comparing 2013 and 2019 estimates) with some areas in Amuru and Adjumani showing increased suitability over time. In contrast, the spatio-temporal model showed that there was a in the median and range of abundance of tsetse in areas where Tiny Targets were deployed. In particular, catches predicted to be high (>10 tsetse/trap/day) were absent in areas where Tiny Targets were deployed.

In 2019, as the national incidence of gHAT declined to record lows, Uganda commenced scale back of tsetse control operations in Maracha district, and currently (January 2024) there are no plans to deploy Tiny Targets in the future. This will mark the first time that no Tiny Targets are deployed in Uganda in over ten years. Our results describe the impact of a successful national tsetse control programme and also produce maps which identify places highly suitable for tsetse and where they may rebound fastest.

Our findings add to earlier smaller-scale studies showing that Tiny Targets are a highly cost-effective method of controlling gHAT vectors [54, 55] leading to their adoption in national programmes to eliminate gHAT [3]. We leverage one of the most data-rich longitudinal datasets of abundance of riverine tsetse in existence (31,553 trapping days), to produce a high-spatial resolution spatio-temporal geostatistical model of tsetse abundance across a ∼16,000km^2^ area (within which Tiny Targets were deployed over an area of ∼4,000km^2^). This approach expanded upon earlier work performed in select districts, and for one time period (2010) [16]. Here, we produce a separate model which considered both spatial and temporal variation, as well as the incorporation and assessment of intervention measures on abundance, through information on the deployment of Tiny Targets between 2011-2019. The spatio-temporal model outperforms that for the previous spatial analysis when looking at metrics of predictive power for known trapping locations, i.e., RMSE of 8.02 flies *vs* 15.2. However, the RMSE for our model remains inflated due to the generation of mean estimates across posterior samples containing high numbers of zeros. Metrics looking at the accuracy of categorised predictions, i.e., low (0-1 flies), medium (>1-10), and high (>10) indicate greater accuracy than direct counts (Appendix: Figure 3).

By performing a counterfactual analysis using the fitted ZINB geostatistical model and varying the intervention category, we show that Tiny Targets reduced the catches of tsetse from monitoring traps between 2011 and 2019. Our results accord with analyses from trials of Tiny Targets in Uganda [7, 8] and suggest that implementation by District tsetse control teams was highly effective. Our results are also comparable to those from the Democratic Republic of Congo where a 85.5% reduction was attributed to the deployment of Tiny Targets [15]. In Chad, an even higher level of control (99.5%) was achieved, [10] probably reflecting local agro-ecological differences. The most important of such differences was perhaps that intervention in Chad was directed against a relatively small and isolated population of tsetse associated with a wetland, whereas in Uganda and DRC the tsetse population was distributed throughout a complex and extensive river network.

Our analyses identified several ‘hotspots’ within the intervention areas where tsetse are predicted to be relatively abundant (Appendix: Figure 4). Indeed, whilst the abundance of tsetse declined over time (Figure 5), we did not see complete elimination of tsetse within areas which have been subject to prolonged control. Tiny Targets are deployed to reduce populations to a level where transmission is interrupted rather than eliminate tsetse themselves. Modelling analysis indicates a ∼60% population suppression is required to achieve interruption of transmission within DRC [56].

The predicted reduction of tsetse in areas where Tiny Targets were not deployed, e.g., central Adjumani and southern Arua (Figure 5), may be attributable to temporal changes not explicitly incorporated within our model but which were captured via the inclusion of a temporal random effect, i.e., noise within the auto-regressive order 1 model, and a linear temporal trend (*RR* = 0.98, 95% *CI* = 0.97-0.98). Additional temporally varying covariates to consider incorporating within future iterations of the model include human population density, climatic variables such as precipitation, and land-use change, which may further explain the temporal trends observed within the data [57, 58]. North-western Uganda has experienced large levels of development within the last decade, primarily due to an influx of refugees resulting in land-use change, i.e., degraded grasslands, woodlands, and tree plantations [59, 60] and human population growth (averaging 3.4% between 2010 and 2020)[61], among other factors. Despite these developments, our species distribution models show that things have either remained the same or improved for tsetse.

Further research is required to determine the link between suitable tsetse habitat and/or tsetse abundance and the geographical distribution of reported gHAT cases. Prior work has shown that proximity to Tiny Targets reduces risk of gHAT in north-western Uganda [16]. However, not all tsetse infested areas are areas of gHAT risk, with tsetse also transmitting trypanosome species pathogenic to livestock but not to humans [19, 62]. Modelling the spatio-temporal variation in gHAT risk requires not only the accurate quantification of the distribution and abundance of parasite, vector, and host populations but also treatment seeking behaviours and diagnostic accessibility. Quantifying each of these factors would aid planning and implementation of interventions to eliminate transmission of gHAT.

Uganda, along with other countries which have eliminated gHAT, or are preparing elimination dossiers for submission to WHO, need to identify and monitor remaining tsetse populations [63]. The methods and predictions described here may be combined with estimates of geographic accessibility to provide a rationale for the placement of cost-effective sentinel monitoring sites to monitor and confirm tsetse population suppression, as demonstrated by Longbottom *et al*. 2020 [64]. Additionally, the models produced identify locations for which we have the least certainty regarding abundance of tsetse, aiding the identification of areas where baseline data may improve our understanding, thus quantifying a process which was previously driven solely by expert opinion and ease of sampling.

## Conclusions

We show that a large-scale national programme of tsetse control, covering ∼4,000 km^2^ across seven districts, in which district-level teams deployed Tiny Targets, greatly reduced the overall abundance of tsetse and contributed to the elimination of gHAT as a public health problem in Uganda.

Tiny Targets reduced the abundance of tsetse in all areas, but tsetse remained at low numbers, particularly in places where the habitat and environment are highly suitable for them. Such sites should be monitored for any re-bound of tsetse and transmission of gHAT. Maps produced by this study can help to optimise surveillance strategies.

There was no clear and consistent decline in the environmental suitability for tsetse, suggesting that natural and anthropogenic change have had little impact on tsetse in north-western Uganda over the last decade.

## Supporting information

Supplementary File - Data for model

