## Supplementary File 1 - Methods supplement for "Impact of a national tsetse control programme to eliminate Gambian sleeping sickness in Uganda: a spatio-temporal modelling study"

### Supplementary Figures

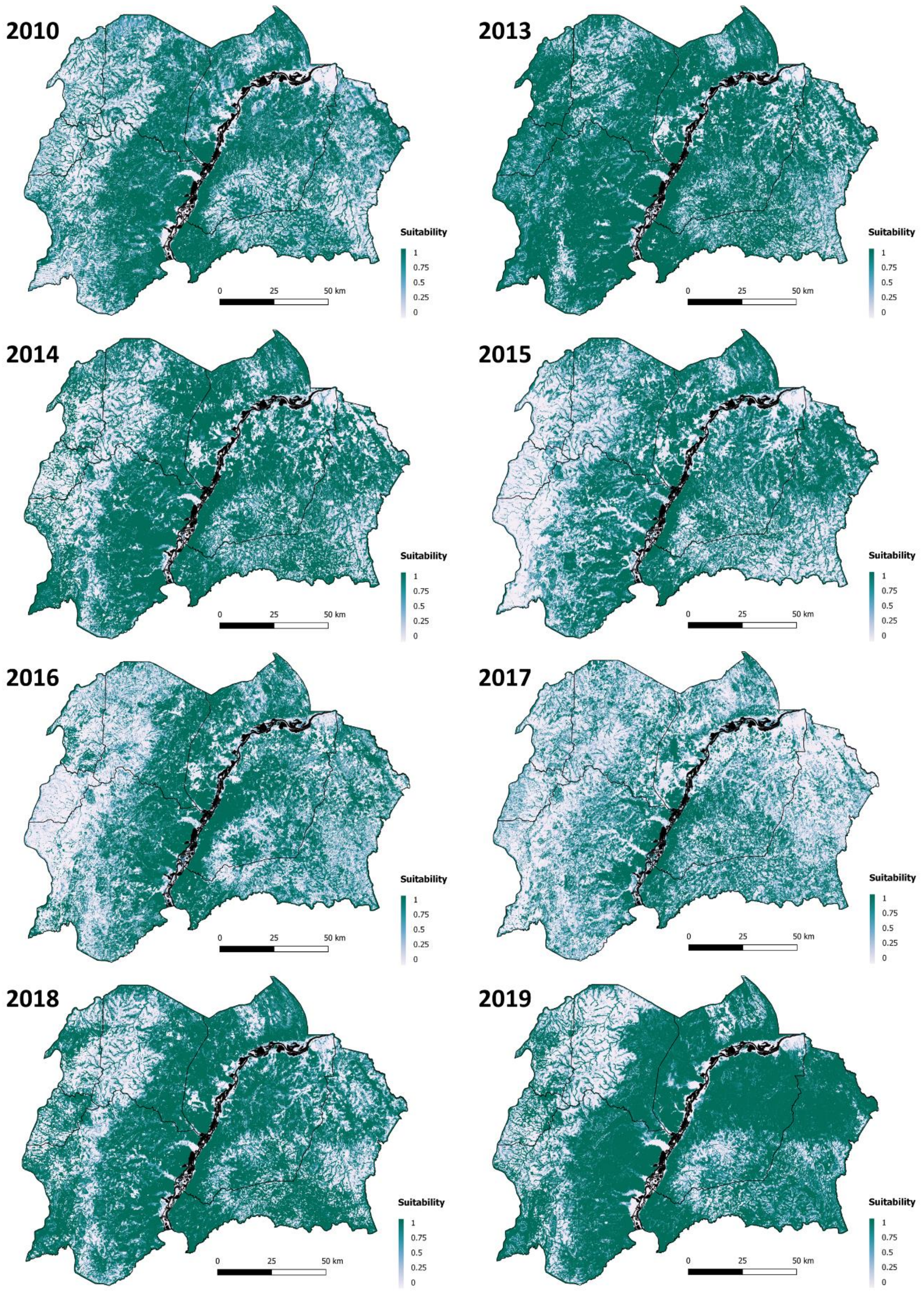

Supplementary Figure 1: Environmental suitability for *G. f. fuscipes* within north-western Uganda, 2010:2019.

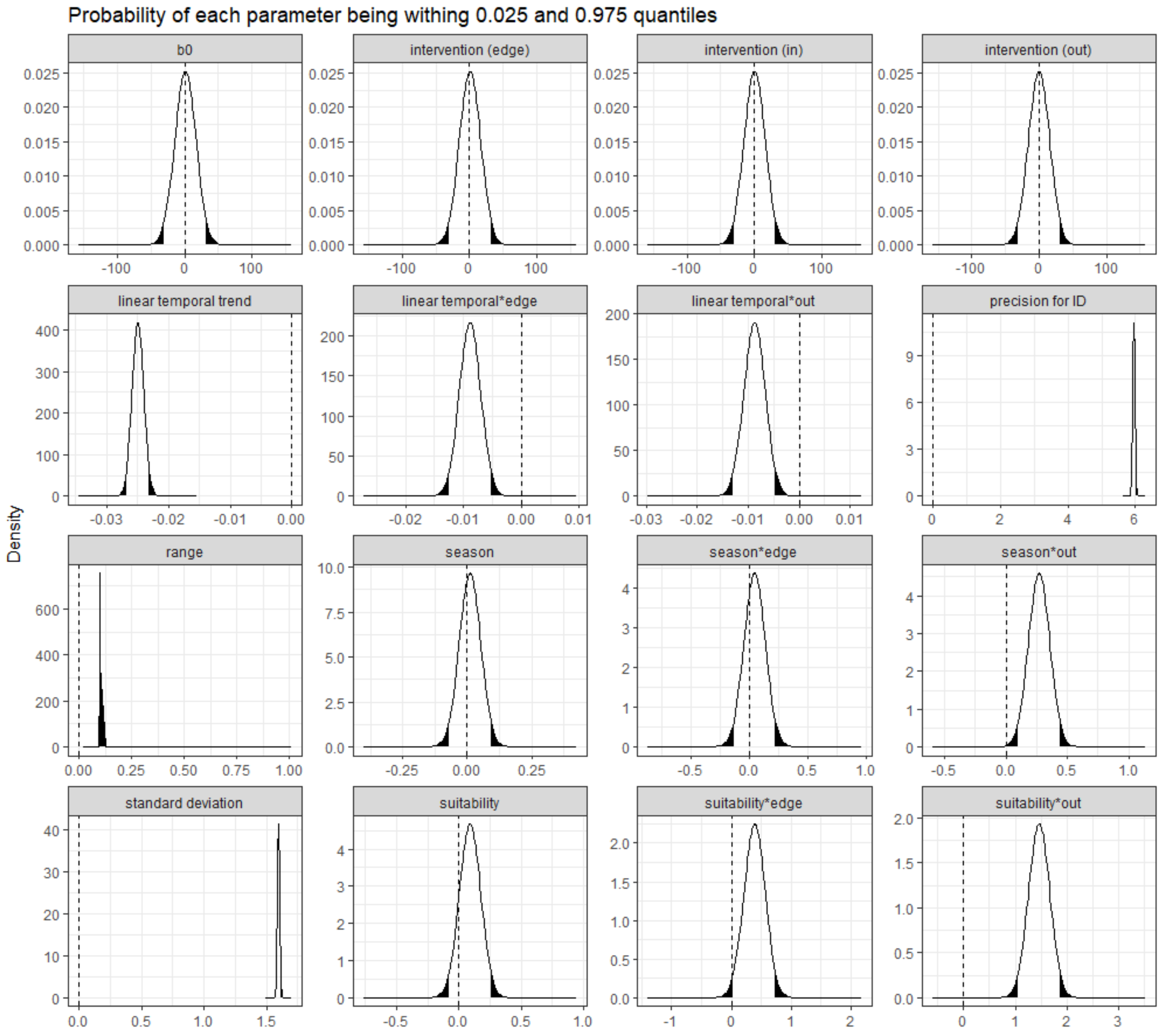

Supplementary Figure 2: Posterior distributions for model parameters, including 2.5% and 97.5% credible intervals (*CrI*).

Black shaded areas represent the 2.5% and 97.5% intervals.

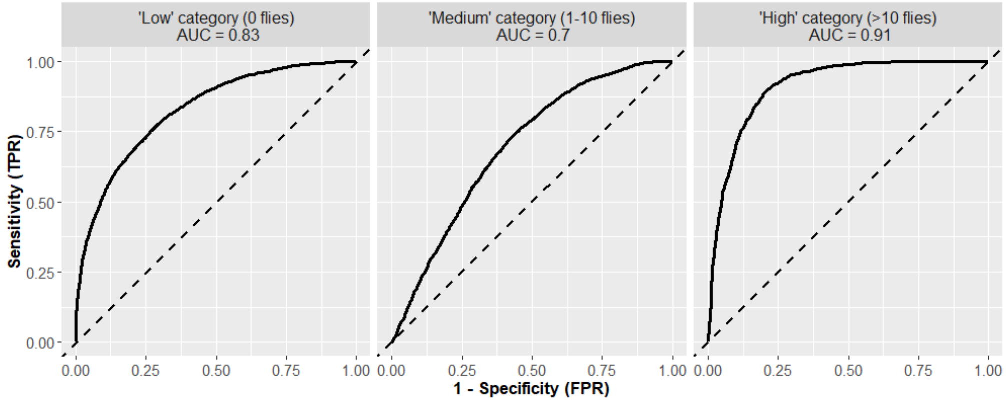

Supplementary Figure 3. Receiver operating characteristic (ROC) curves for prediction categories.

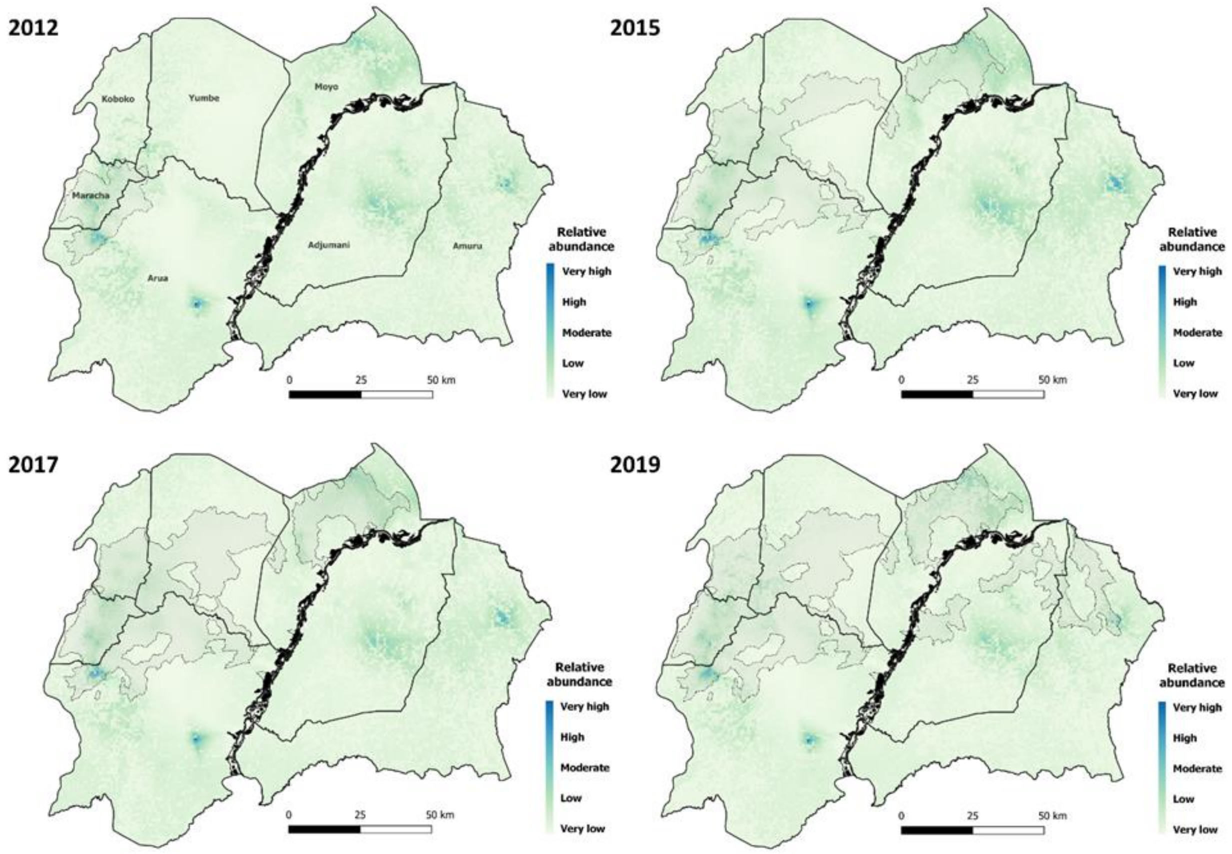

Supplementary Figure 4. Comparison of relative abundance of *G. f. fuscipes* abundance during four time periods.

The prediction period relates to February of each year (2012, 2015, 2017 and 2019). Abundance categories are based off quantiles (values for each year presented as Supplementary Table 8). Grey dotted polygons represent the watersheds covered by the TT intervention during February of the corresponding year, watershed data obtained from [1].

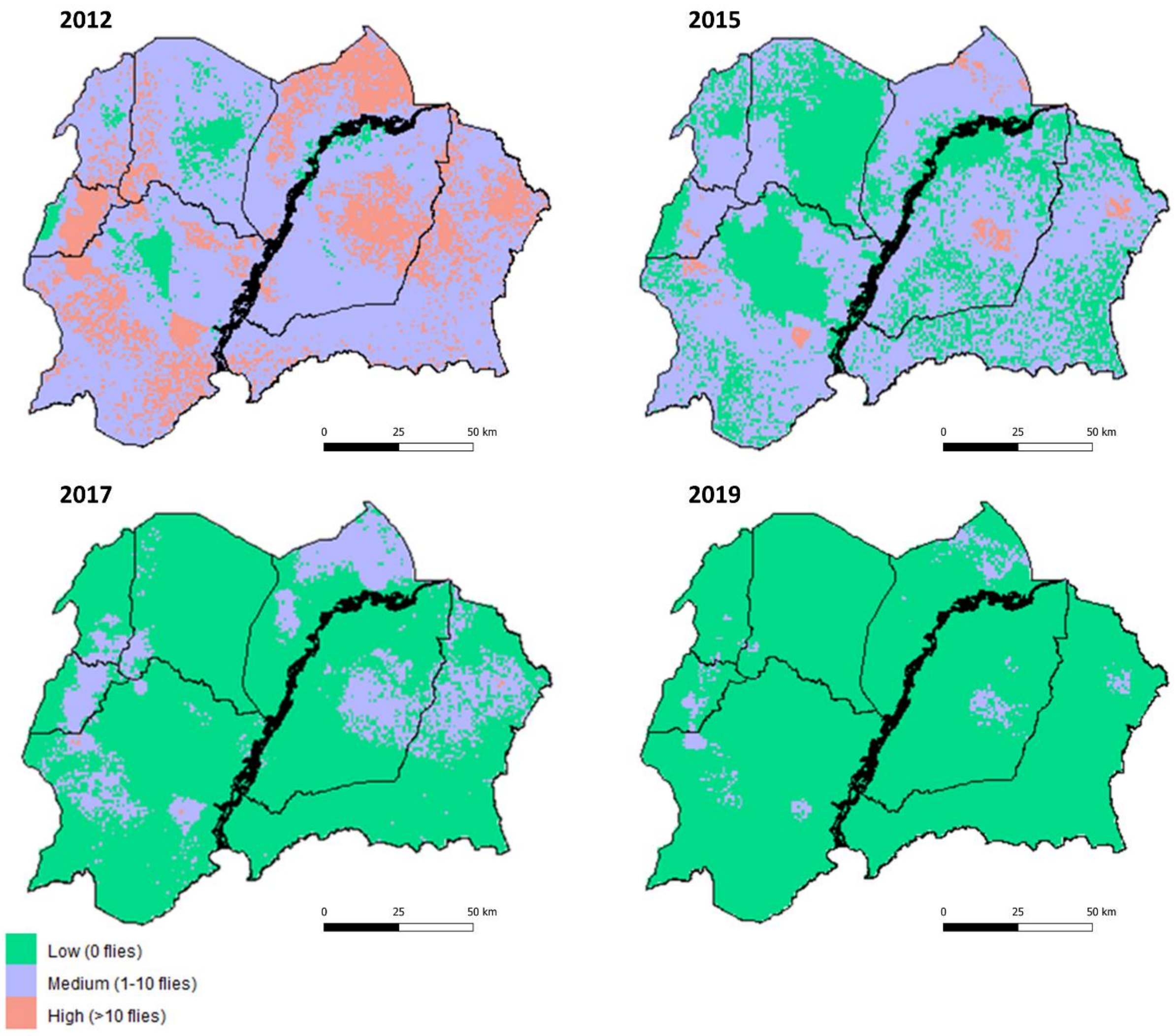

Supplementary Figure 5. Categorised predictions of the number of tsetse per cell.
Results are produced for February of each year and represent the mean number of flies per trapping day.

### Supplementary Tables

#### Supplementary Table 1. Tiny Target deployment data throughout north-western Uganda.

| *Deployment date* | *Adjumani* | *Amuru* | *Arua* | *Koboko* | *Maracha* | *Moyo* | *Yumbe* | *Number of targets distributed* |
|---|---|---|---|---|---|---|---|---|
| *Nov 2011-Dec 2012* | | | X | | X | | X | 1,336 |
| *Dec 2012 - Dec2013* | | | X | | X | | X | 2,872 |
| *Jan – Mar 2014* | | | X | | X | X | X | 6,850* |
| *Nov 2014 - Feb 2015* | | | X | X | X | X | X | 17,419 |
| *Apr - May 2015* | | | X | X | X | X | X | 21,050 |
| *Jan 2016* | | | X | X | X | X | X | 22,026 |
| *Jun - Jul 2016* | | | X | X | X | X | X | 14,826 |
| *Dec 2016 - Mar 2017* | | | X | X | X | X | X | 16,821 |
| *Jun - Jul 2017* | X | X | X | X | X | X | X | 19,355 |
| *Dec 2017 - Feb 2018* | X | X | X | X | X | X | X | 17,402 |
| *Jun - Aug 2018* | X | X | X | X | X | X | X | 17,400* |
| *Jan 2019* | X | X | X | X | X | X | X | 20,205 |
| *Mid 2019* | X | X | X | X | X | X | X | 22,216 |

**Estimated numbers*

#### Supplementary Table 2. Variables collected at each trapping location.

| *Variable* | *Description* |
|---|---|
| *SITEID* | Unique identifier for the site in which sampling was performed. |
| *NORTHING* | The northward-measured y-coordinate of the trap location (recorded in the UTM 36N coordinate reference system). |
| *EASTING* | The eastward-measured x-coordinate of the trap location (recorded in the UTM 36N coordinate reference system). |
| *DISTRICT* | The district in which the trap was situated. |
| *RIVER* | The name of the river along which the trap was deployed (if applicable). |
| *DATE_SETUP* | The date of deployment for the trap (recorded in serialized date format). |
| *DATE_COLLECTED* | The date of collection for the trap (recorded in serialized date format, traps visited after 24 and 48 hours). |
| *MONTH* | The month in which the trap was deployed. |
| *YEAR* | The year in which the trap was deployed. |
| *SPECIES* | The species of tsetse obtained through sampling at that location. |
| *MALES* | The number of male tsetse obtained through sampling at that location. |
| *FEMALES* | The number of female tsetse obtained through sampling at that location. |
| *UNKNOWN* | The number of tsetse obtained through sampling at that location for which a sex identification was not possible. |
| *TOTAL* | The total number of tsetse obtained through sampling at that location (sum of males, females and unknown). |

#### Supplementary Table 3. Summaries of the collated *G. f. fuscipes* data, given per district.

Summaries of mean flies/trap/day consist of traps inside and outside of intervention areas and are one mean generated for the whole sample period.

| *District* | *Sampled sites* | *Flies caught* | *Days deployed* | *Mean flies/trap/day* |
|---|---|---|---|---|
| *Adjumani* | 91 | 484 | 426 | 1.14 |
| *Amuru* | 22 | 202 | 204 | 0.99 |
| *Arua* | 219 | 22517 | 14269 | 1.58 |
| *Koboko* | 75 | 10613 | 5589 | 1.90 |
| *Maracha* | 82 | 16107 | 7735 | 2.08 |
| *Moyo* | 41 | 833 | 815 | 1.02 |
| *Yumbe* | 39 | 1788 | 2515 | 0.71 |
| ***Total*** | **569** | **52544** | **31553** | **1.67** |

#### Supplementary Table 4: Number of presence and absence records available per year.

Records relate to traps outside of an intervention area only.

| Year | Presence | Absence |
|---|---|---|
| 2010 | 102 | 51 |
| 2011 | 50 | 3 |
| 2012 | 22 | 0 |
| 2013 | 28 | 1 |

| | | |
|---|---|---|
| 2014 | 15 | 3 |
| 2015 | 18 | 3 |
| 2016 | 42 | 17 |
| 2017 | 69 | 77 |
| 2018 | 12 | 4 |
| 2019 | 18 | 3 |

#### Supplementary Table 5: The relative contribution of each covariate to the boosted regression tree model.

| Covariate | Relative influence (%) |
|---|---|
| Elevation | 19.14 |
| Normalised difference vegetation index (NDVI) | 18.57 |
| Distance to rivers | 18.48 |
| Land surface temperature (LST) | 16.28 |
| Proportion of vegetation (PVI) | 15.86 |
| Slope (% gain) | 11.67 |

#### Supplementary Table 6. Summary of model evaluation statistics for the varying spatio-temporal models explored.

| Model | Fixed effects: Additive fixed effects | Fixed effects: Interaction term | Random effect | DIC | WAIC | CPO |
|---|---|---|---|---|---|---|
| 1 | | | Site ID | 37064.59 | 37128.27 | 19710.57 |
| 2 | Intercept, Suitability | | Site ID | 37008.64 | 37066.98 | 19493.13 |
| 3 | Intercept, Suitability, Categorised intervention | | Site ID | 36661.47 | 36713.62 | 19111.56 |
| 4 | Intercept | Suitability*Categorised intervention | Site ID | 36618.51 | 36683.19 | 18757.23 |
| 5 | Intercept | Suitability*Categorised intervention<br>Season* Categorised intervention | Site ID | 36599.59 | 36657.30 | 18831.86 |
| 6 | Intercept | Suitability*Categorised intervention<br>Season* Categorised intervention<br>Linear temporal trend*Categorised intervention | Site ID | 35547.63 | 35593.69 | 17908.93 |
| 7 | Intercept | Suitability*Categorised intervention<br>Season* Categorised intervention<br>Linear temporal trend*Categorised intervention | | 35556.95 | 35605.43 | 17909.37 |
| 8 | Intercept | Suitability*Categorised intervention<br>Season* Categorised intervention | | 35590.95 | 35632.27 | 18154.46 |
| 9 | Intercept, Suitability | Suitability*Categorised intervention<br>Season* Categorised intervention | | 35601.33 | 35659.7 | 17870.5 |
| 10 | Intercept, Suitability, Categorised intervention, season, Linear temporal trend | Suitability*Categorised intervention<br>Season* Categorised intervention<br>Linear temporal trend*Categorised intervention | Site ID | 35538.97 | 35583.8 | 17899.82 |

#### Supplementary Table 7: Parameter/hyperparameter values for priors within the spatio-temporal model.

| Function | Prior | Parameter/hyper-parameter | Value |
|---|---|---|---|
| Matérn SPDE model | Penalized Complexity (PC) prior | Spatial range of the random field: $\rho$<br>Marginal standard deviation of the field: $\sigma^2$ | 0.0115, 0.1<br>$\sqrt{1.792362}$, 0.05 |
| $AR_1$ model | LogGamma<br>Normal | Log precision: $\theta_1$<br>Logit lag one correlation: $\theta_2$ (also known as $\rho$) | Initial = 0.75, fixed<br>Default: 0, 0.15 |

#### Supplementary Table 8: Quantile values for the relative abundance points used in Supplementary Figure 4.

| Year | Relative abundance (quantiles): Very Low | Low | Moderate | High | Very high |
|---|---|---|---|---|---|
| 2019 | 0 | 2.5 | 5 | 7.5 | 10 |
| 2017 | 0 | 5.5 | 11 | 16.5 | 22 |

| | | | | | |
|---|---|---|---|---|---|
| 2015 | 0 | 11 | 22 | 33 | 44 |
| 2012 | 0 | 38.75 | 77.5 | 116.5 | 155 |

### Supplementary Methods

#### Sample collection

A preliminary survey to establish fly distribution and density was performed within Arua and Maracha districts during 2010 [2, 3]. Following this, tsetse control began in five blocks within Arua and Maracha between November 2011 and December 2012 [2], expanding to 500km$^2$ within these districts between December 2012 and December 2013 [2, 3]. In January 2014, control coverage expanded to ~3500km$^2$ within Arua, Koboko, Yumbe and Moyo, with a final expansion to g-HAT affected areas within Adjumani and Amuru districts in June 2017 [1]. As of 2019, the highest risk areas within all seven endemic districts were controlled, totalling an area of ~5000km$^2$ [1, 4]. With each geographical expansion, new baseline surveys were performed to ascertain where best to focus control efforts, and longitudinal monitoring sites were established.

#### Data preparation

***Dataset one:*** To ensure that we were modelling suitability in the absence of tsetse control, only data collected between 2010-2019 from traps located ≥5km outside of the Tiny Target intervention area were used. Data was split using an *'intervention'* variable, retaining trapping records situated outside of an intervention area. A binary presence (1) absence (0) indicator variable was produced using the total number of flies caught during each 24-hour sampling period. Trapping records were spatially and temporally aggregated to produce presence/absence data, such that for each unique sampling location, as defined by a unique 30m × 30m grid cell, only one record was retained per calendar year (January – December). If, for the same grid cell, both presence and absence were recorded within the same year, the location was deemed a presence location, and absence records for that location and year were removed. This ensured that no ambiguity occurred within the model when determining suitability at each gridded location.

***Dataset two:*** The raw occurrence dataset, with the addition of the binary and continuous intervention variables, was used for the spatio-temporal model. No spatial-temporal aggregation was performed, with each row in the dataset being representative of a unique 24-hour sampling period and site.

#### Covariate generation process

Elevation (expressed as meters above sea level) was obtained from the Shuttle Radar Topography Mission [5]. Utilising the elevation surface and hydrological tools in ArcMAP 10.4, a flow accumulation surface was generated from the imagery to derive a river network. An accumulation value of 3000 was selected for the stream order, this defined the detail of the constructed network. Using this river network, the Euclidean distance from the network to each 30 × 30m pixel within the study extent was calculated, this produced a 'distance to rivers' covariate, where cell values represented distance in meters. Additionally, utilising the SRTM data, a slope surface was calculated using the 'Slope' tool in the 'Spatial Analyst' toolbox within ArcMAP, this was represented as percentage gain across the study extent. Utilising Landsat 5 and Landsat 8 imagery [6, 7], land-surface temperature (LST, °C), proportion of vegetation (PVI, range = 0-1) and a normalised difference vegetation index (NDVI, range = -1-1) were calculated in R, following the steps outlined by the United States Geological Survey [8]. Synoptic surfaces were generated for elevation, slope and distance to rivers, and annual surfaces were created for LST, PVI and NDVI, for each year between 2011 and 2019.

#### Species distribution model

To produce estimates of *G. f. fuscipes* densities across north-western Uganda in locations for which no sampling was performed, we first generated estimates of habitat suitability utilising a presence-absence SDM framework. SDMs are used to provide an understanding of and/or to predict the distribution of a species across a landscape [9], combining information on species occurrence, i.e., presence or absence at a location, with information on environmental or socio-economic variables (covariates) at the same location [10]. Using a temporally aggregated presence-absence dataset containing one observation per grid cell per year, i.e., if tsetse were caught within a sampled cell at any period within a year, the cell was assigned 'presence' for that year we constructed a presence-absence boosted regression tree (BRT) model using the '*caret*' package within R (version 3.5.1) [11, 12].

To prepare the presence-absence data for use in the model, covariate values were assigned to each observation for the corresponding sampling year. For example, a trapping record obtained during 2015 was allocated a mean covariate value for NDVI, LST and PVI which was deduced from available cloud-free satellite imagery obtained for the first dry season (December-February) of same calendar year, and synoptic covariate values for slope, distance to rivers and elevation. This was the case for most observations except those during 2010, which were assigned 2011 covariate data, and 2012, which were assigned 2013 data, due to the scan line corrector error associated with Landsat 7 preventing clear imagery from being obtained for those years [13]. Following assignment of covariate values, records were checked to ensure that each observation contained a value for each of the six covariates.

The data were partitioned into 'training' and 'test' datasets, with 70% of records being used for model training and 30% withheld for testing purposes. We first employed Bayesian parameter optimization to select values for three tuning parameters needed to implement the BRT method: number of iterations (trees), complexity of the tree (interaction depth) and learning rate i.e., how quickly the algorithm adapts, also referred to as shrinkage [14]. During parameter optimization, the candidate models were evaluated by repeated (three times), 10-fold cross-validation. The cross-validation process separates the data set into *k* subsets containing approximately the same number of occurrence and background points [14, 15]. The sub-model is then iteratively trained using *k-1* data subsets, and the performance in predicting the withheld data is evaluated by the receiver operating characteristic (ROC) [16]. The 'optimal' model across these parameters, as defined by ROC, was chosen, and the parameters from this model were then used to fit the model to the partitioned training dataset.

The optimal model performance against the 30% test dataset was evaluated by both the Brier score [17], and area under curve (AUC) values [18]. The Brier score measures the difference between observed and fitted values, where values are assigned 1 if tsetse were observed/predicted to be present, and 0 otherwise [17, 19]. More specifically, the Brier score is defined as $\frac{1}{n}\sum_{i=1}^{n}(y_i - \hat{p}_i)^2$ where $y_i = 1$ if flies were present at location $i$, and zero otherwise, and $\hat{\mathrm{p}}_\mathrm{i}$ is the cross-validated predicted probability that flies were present. As such, the Brier score lies between 0 (perfect predictions) and 1. As the Brier score is evaluated at known locations, it can be used to visualise spatial errors and interrogate uncertainty. The AUC represents the area under the ROC curve, where the ROC curve is a plot of true positive rate *vs* false positive rate at different classification thresholds [16]. An AUC value of 0.5 is equivalent to a 'random draw' prediction, with values ≥0.7 indicating a good model fit. The fitted model was then used with covariates for each year between 2011-2019 to produce annual estimates of suitability across north-western Uganda at a 30m × 30m resolution, as informed by the observed data.

#### Geostatistical model description

The underlying statistical model was a spatially and temporally explicit hierarchical generalised linear regression model for ZINB data, using the log link function and a type 1 likelihood. A type 1 likelihood accounts for two different types of zeros within the dataset: *structural* or true zeros which represent the true absence of the species in a location, and *sampling* zeros, where a zero is recorded as a reflection of sampling effort/chance, i.e., a species may be present at that location however was undetected [20]. Suppose that structural zeros occur with probability $\pi$ and sampling zeros occur with probability $1 - \pi$. Therefore, the probability distribution of the ZINB random variable $y_i$ can be written:

$$\Pr(y_{ij} = n) = \begin{cases} \pi_{ij} & \text{if } n = 0 \\ (1 - \pi_{ij}) g(y_{ij}) & \text{if } n > 0 \end{cases}$$

Where $g(y_i)$ is the negative binomial distribution. When accounting for both space $(s)$, i.e., each 30m x 30m pixel across north-western Uganda, and time $(t)$, i.e., each month between October 2010 and December 2019, the data model utilising the negative binomial distribution can be written as:

$$Y(s_i, t_j) \sim NegBin(\mu(s_i, t_j) = d(s_i, t_j)\, \eta(s_i, t_j), \alpha)$$

Where $Y(s_i, t_j)$ is the number of flies caught in trap $i$, at location $s$, in month $j$. $\mu(s_i, t_j)$ is the mean number of flies observed at trap $i$ in month $j$, $\eta(s_i, t_j)$ is the rate of flies caught per trap day, the offset $d(s_i, t_j)$ represents the number of days over which flies were caught at trap $i$, during month $j$, and $\alpha$ represents the overdispersion parameter.

The process model can then be defined as:

$$\log\left(\mu(s_i, t_j)\right) = \log\left(d(s_i, t_j)\right) + \log\left(\eta(s_i, t_j)\right)$$

$$\log\left(\eta(s_i, t_j)\right) = \beta_0 + \boldsymbol{\beta^T X}(s_i, t_j) + U(s_i, t_j) + V(s_i, t_j)$$

$$\log(\mu(s_i, t_j) = \beta_0 + \boldsymbol{\beta^T X}(s_i, t_j) + \log\left(d(s_i, t_j)\right) + U(s_i, t_j) + V(s_i, t_j)$$

$$U(s_i, t_j) \sim N(0, \sigma^2)$$

$$V(s_i, t_j) = W(s_i)\, Z(t_j)$$

$$W(s) \sim SGP(O, \textstyle\sum),$$

$$Z(t) \sim AR_1(t)$$

Where the coefficient $\beta_0$ represents the intercept, $X(s_i, t_j)$ are covariates associated with trap $i$ observed at month $j$ with $\beta^T, T = 1, \ldots, n$ representing the associated coefficients. $U(s_i, t_j)$ represents the spatially uncorrelated random effects, zero-mean Gaussian random variables, and $V(s_i, t_j)$ represents the spatially and temporally structured random effects, which consists of $W(s)$, a zero-mean stationary Gaussian process with a Matérn covariance function, and $Z(t)$, defined by the covariance function corresponding to a discrete-time autoregressive stochastic process of the first order ($AR_1$). We construct a separable spatio-temporal model, where the correlation between $W(s)$ and $Z(t)$ is separate and additive.

#### Model fitting and validation

Calculating the predictive error of a model is commonly performed using a cross-validation approach, in which data are repeatedly split into two subsets, one used for model training and the other for model testing [21]. When considering spatial autocorrelation, attempts should be made to ensure that data used within training and testing datasets are independent [22]. If training and testing data are close in space, but predictions are being made in locations far from the training data, there is a risk of under-estimating predictive error [23]. A common cross-validation method for spatio-temporal data, which accounts for spatial autocorrelation and resolves under-estimation of predictive error, involves leave-*n*-out cross-validation approaches which incorporate a distance-based buffer around hold-out points [21, 24]. This approach ensures that *n* randomly selected points, and those correlated with it (specified by a buffer exceeding the range of spatial autocorrelation) are removed from the training dataset and are used to evaluate model performance. Within our work, model validation was performed using a spatial leave-one-out cross-validation (SLOO-CV) approach, based on an adaptation of methods described in [23, 24]. Namely, the steps performed are:

1. Remove one spatial location and all corresponding temporal observations from the initial dataset (*n* = 416 locations).
2. Remove observations within a radius of the selected location, where the radius is a value exceeding the range of the spatial autocorrelation, obtained from the GMRF.
3. Predict tsetse abundance at the location of the removed observation using parameters estimated using the remaining data within the initial dataset.

Steps (1) to (3) are repeated $k$ times. $k$ can be less than or equal to the number of data points, $n$, with $k = n$ being the case where all data points are left out once [23]. Validation statistics were then generated for the returned predictions, which included assessing the correlation between the predicted and observed tsetse densities through generating summaries of the root-mean-square error (RMSE), as well as the mean absolute error (MAE) [25], across the posterior predictions.

radar-topography-mission-srtm-1-arc?qt-science_center_objects=0#qt-science_center_objects.
6. U.S. Geological Survey, *Landsat-5 imagery courtesy of the U.S. Geological Survey.*
7. U.S. Geological Survey, *Landsat-8 imagery courtesy of the U.S. Geological Survey.*
8. U.S. Geological Survey. *Using the USGS Landsat Level-1 Data Product*. 2020 8th July 2020]; Available from: https://www.usgs.gov/land-resources/nli/landsat/using-usgs-landsat-level-1-data-product.
9. Elith, J. and J.R. Leathwick, *Species Distribution Models: Ecological Explanation and Prediction Across Space and Time.* Annual Review of Ecology, Evolution, and Systematics, 2009. **40**(1): p. 677-697.
10. Barry, S. and J. Elith, *Error and uncertainty in habitat models.* Journal of Applied Ecology, 2006. **43**(3): p. 413-423.
11. Kuhn, M., *caret: Classification and Regression Training. R package version 6.0-86*. 2020.
12. R Core Team, *R version 3.5.1 (2018-07-02) -- "Feather Spray"*. 2020.
13. U.S. Geological Survey. *Landsat Missions: Landsat 7*. 2021 26th April 2021]; Available from: https://www.usgs.gov/core-science-systems/nli/landsat/landsat-7?qt-science_support_page_related_con=0#qt-science_support_page_related_con.
14. Elith, J., J.R. Leathwick, and T. Hastie, *A working guide to boosted regression trees.* Journal of Animal Ecology, 2008. **77**(4): p. 802-813.
15. Stone, M., *Cross-Validatory Choice and Assessment of Statistical Predictions.* Journal of the Royal Statistical Society. Series B (Methodological), 1974. **36**(2): p. 111-147.
16. Zou, K.H., A.J. O'Malley, and L. Mauri, *Receiver-Operating Characteristic Analysis for Evaluating Diagnostic Tests and Predictive Models.* Circulation, 2007. **115**(5): p. 654-657.
17. Brier, G.W., *Verification of forecasts expressed in terms of probability.* Monthly Weather Review, 1950. **78**(1): p. 1-3.
18. DeLong, E.R., D.M. DeLong, and D.L. Clarke-Pearson, *Comparing the Areas under Two or More Correlated Receiver Operating Characteristic Curves: A Nonparametric Approach.* Biometrics, 1988. **44**(3): p. 837-845.
19. Tirados, I., et al., *Impact of tiny targets on Glossina fuscipes quanzensis, the primary vector of human African trypanosomiasis in the Democratic Republic of the Congo.* PLOS Neglected Tropical Diseases, 2020. **14**(10): p. e0008270.
20. Arab, A., *Spatial and Spatio-Temporal Models for Modeling Epidemiological Data with Excess Zeros.* International journal of environmental research and public health, 2015. **12**(9): p. 10536-10548.
21. Roberts, D.R., et al., *Cross-validation strategies for data with temporal, spatial, hierarchical, or phylogenetic structure.* Ecography, 2017. **40**(8): p. 913-929.
22. Araújo, M.B., et al., *Validation of species–climate impact models under climate change.* Global Change Biology, 2005. **11**(9): p. 1504-1513.
23. Lucas, T., A. Python, and D. Redding, *Graphical outputs and Spatial Cross-validation for the R-INLA package using INLAutils.* arXiv preprint arXiv:2004.02324, 2020.
24. Le Rest, K., et al., *Spatial leave-one-out cross-validation for variable selection in the presence of spatial autocorrelation.* Global Ecology and Biogeography, 2014. **23**(7): p. 811-820.
25. Chai, T. and R.R. Draxler, *Root mean square error (RMSE) or mean absolute error (MAE)? – Arguments against avoiding RMSE in the literature.* Geosci. Model Dev., 2014. **7**(3): p. 1247-1250.
